# Evolutionary Strategies Applied to Artificial Gene Regulatory Networks

**DOI:** 10.1101/2021.09.28.462218

**Authors:** André L. L. Moreira, César Rennó-Costa

## Abstract

Evolution optimizes cellular behavior throughout sequential generations by selecting the successful individual cells in a given context. As gene regulatory networks (GRNs) determine the behavior of single cells by ruling the activation of different processes - such as cell differentiation and death - how GRNs change from one generation to the other might have a relevant impact on the course of evolution. It is not clear, however, which mechanisms that affect GRNs effectively favor evolution and how. Here, we use a population of computational robotic models controlled by artificial gene regulatory networks (AGRNs) to evaluate the impact of different genetic modification strategies in the course of evolution. The virtual agent senses the ambient and acts on it as a bacteria in different phototaxis-like tasks - orientation to light, phototaxis, and phototaxis with obstacles. We studied how the strategies of gradual and abrupt changes on the AGRNs impact evolution considering multiple levels of task complexity. The results indicated that a gradual increase in the complexity of the performed tasks is beneficial for the evolution of the model. Furthermore, we have seen that larger gene regulatory networks are needed for more complex tasks, with single-gene duplication being an excellent evolutionary strategy for growing these networks, as opposed to full-genome duplication. Studying how GRNs evolved in a biological environment allows us to improve the computational models produced and provide insights into aspects and events that influenced the development of life on earth.

## 1. INTRODUCTION

*Natural selection* is a process in which individuals with distinct heritable characteristics are subjected to selective pressure, resulting in the survival of those more fitted to that environment. The reproductive success of those that survive allows the passage of their alleles to the next generation (REECE et al., 2015, p. 488). Four conditions are necessary for natural selection to occur (RIDLEY, 2006): (1) reproduction: individuals reproduce to form a new generation; (2) heredity: subsequent generations have characteristics inherited from their parents; (3) variation between individual characters: individuals of the same generation have different characteristics, and (4) aptitude variation: different individuals perform differently in the different tasks, allowing the definition of more or less adapted beings. Natural selection is the core of what we know as evolution, and, consequently, the specifics of how each of these processes is implemented strongly impact how evolution develops.

Computer simulations have been widely used to study the dynamics of evolution, as, for example, the evolutionary programming technique proposed by Fogel (1962). These models implement relevant aspects of a behaving agent with specific rules for selection and reproduction. For example, bacteria are represented through the modeling of Gene Regulation Networks (GRNs), a strong abstraction of biological systems (HUYNH-THU; SANGUINETTI, 2019), and play an important role in the translation between spatial information patterns present in the environment and dynamic processes in time. In other words, gene regulation networks influence the behavioral response of individuals when subjected to different contexts (CUSSAT-BLANC; HARRINGTON; BANZHAF, 2019). In addition, through the regulation of protein production, GRNs influence essential processes in life, such as cell differentiation (DAVIDSON et al., 2002, GÉRARD; TYS; LEMAIGRE, 2017), metabolism (WATSON; WALHOUT, 2014), and evolution ( DAVIDSON, 2010). GRN models allow the study of both the behavioral and the morphological evolution of an animal since these characteristics are the product of its development process, which is encoded in its species regulatory genome. Thus, morphological changes in a species result from the change in its gene regulation (DAVIDSON, 2010).

Here we use the Artificial Gene Regulatory Network (AGRN) model proposed by Banzhaf (2003) to study the impact of different strategies for gene replication on the course of evolution. In the model, genes and their interactions are translated into graphs (Figure 1). By classifying Nodes (genes) into sensors, processors, and controllers, we simulated the interaction between network and environment through sensor genes and the behavior modulation through controller genes. The critical question is, which is the preferred mechanism for varying an AGRN to support better results in an evolutionary process? In this sense, this work aims to evaluate how different evolutionary strategies, such as changes in the species’ context (change of habitat, variation in the complexity of the tasks) and individuals’ physiological changes, impact the evolutionary process.

**Figure 1:**
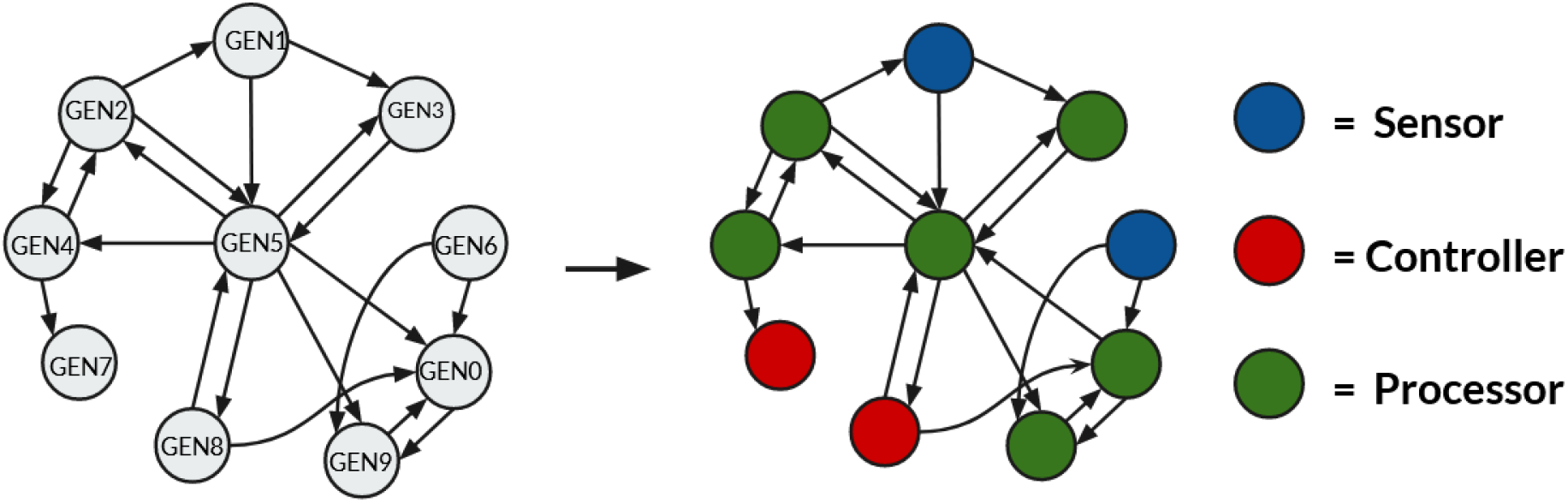
Translation of a hypothetical gene regulation network into the artificial gene regulation network model. Nodes represent genes of the classes: sensor, processor, and controller. Vertices represent excitatory and inhibitory relationships between genes.

## 2. METHODOLOGY

### 2.1 Body Model

The structure of each simulated agent was based on a generic bacteria with the ability to move around in an environment, identify objectives to be captured, and avoid obstacles during tasks. For this, we base the behaving model on the Epuck robot (MONDADA et al., 2009) (Figure 2). The simulated agent has proximity sensors distributed around its contour. The model includes a total of seven proximity sensors. The frontal sensor can differentiate objectives and obstacles in the environment. In addition, four controllers that define behaviors were simulated, namely: accelerate forward, accelerate backward, rotate right, and rotate left

**Figure 2:**
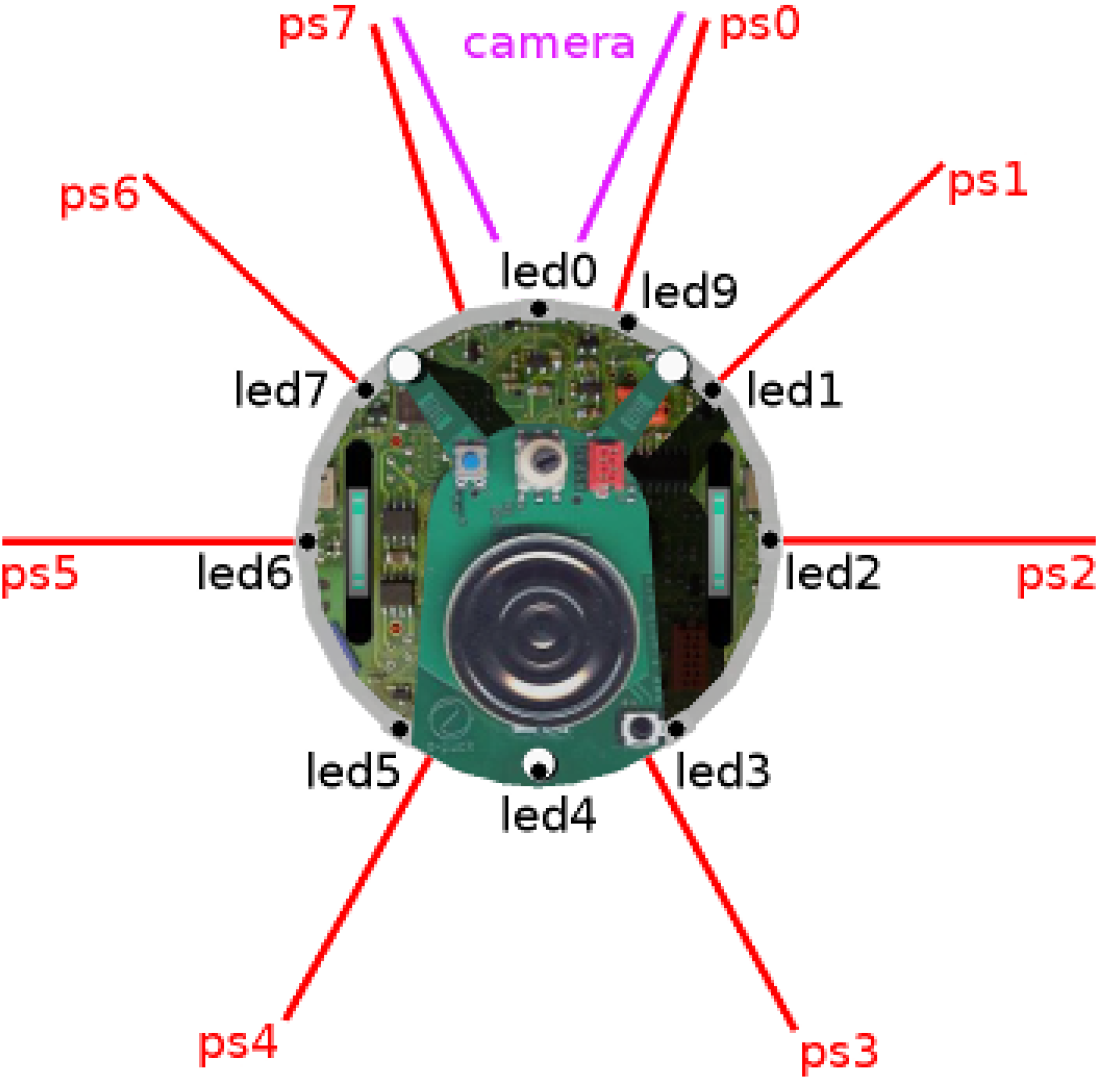
Epuck Structure. Consisting of 8 proximity sensors, a front camera and two wheels capable of turning in both directions. **Source:** Epuck Documentation by the webbots program^1^

### 2.2 The Artificial Gene Regulation Network Model

We use an extension of the Artificial Gene Regulation Network (AGRN) model (Figure 3), proposed by Banzhaf (2003), composed of genes (nodes), which assume different expression levels and are interconnected, creating gene-protein interactions (edges). Three gene classes make up our model: sensors, processors, and controllers. Sensors interact directly with the processing layer and receive input from various environmental information: position, object proximity, and differentiation. In addition to receiving input from sensors, processor genes interact, creating recurrence relationships and modulating the controller class genes. The controller genes define the individual’s actions, which activate the functions: accelerating, braking, and rotating.

**Figure 3:**
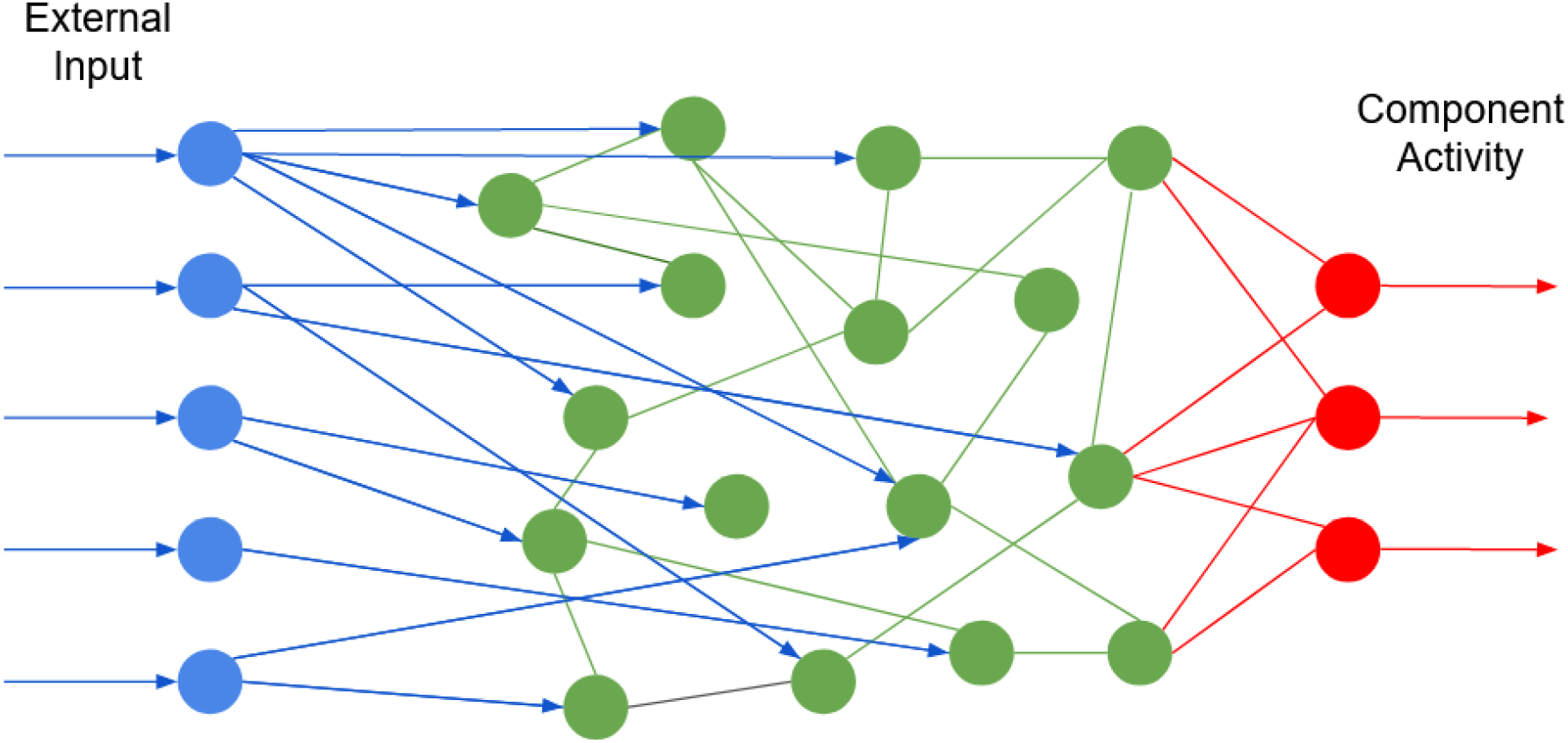
Network with nodes representing genes. In blue, sensor genes that receive input from the environment. In green, processor genes. In red, controllers that activate and deactivate individual functions. Edges represent interactions between genes, which can be inhibitory or excitatory. Green (between processor genes) and red (between controllers and processors) edges can be bidirectional or not. The blue ones (between sensors and processors) are unidirectional.

Each agent is represented by a graph that keeps pairwise interactions between each of the simulated genes. During a simulation, the structure of sensor and controller genes, which represent the bacterium’s physical structure, is defined by the task they are performing. In turn, the amount of processor genes is a study parameter. The random creation of an individual consists of generating random relationships between the genes in the network, with −1 being the minimum inhibition relationship, and 1 the maximum excitation between two genes.

The behavior of processor and controller genes is based on a Perceptron (NIELSEN, 2015): their individual activity is given by a function that relates all inputs received from other genes, both positive and negative. However, unlike a common Perceptron, they do not go through an activation function with binary output (active or inactive). In this investigation, each gene’s activity is represented by a real number, which can vary between 0 and 1, translating the different gene expression levels observed biologically. At each moment of the simulation, the activity of each gene is recalculated, and its new activity is defined by the following equation:

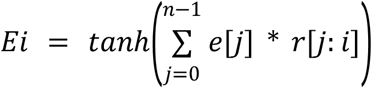

In this equation, “E” is the energy of the calculated gene “i”, “e” is the energy of the genes ‘j’ that are acting on it, and “r” is the excitation or inhibition relationship between the genes.

Sensor genes have their activity modulated by their function. Proximity sensors activate when there is a light (objective to be captured) in their activation area, the sensor sensor activation energy is defined by the distance from the identified target: its energy is 1 when the target is very close and it gets weaker the farther away it is, tending to 0 over very large distances. The front sensor identifies obstacles or threats, being 1 when an obstacle is in its operation area and 0 when there are no threats ahead.

Controller genes, like others, have their energy varying between 0 and 1 during simulations. However, in addition to interfering with the network by exciting and inhibiting other genes, controller genes interfere in the simulated individuals’ behavior. The intensity with which these genes act in the robot is related to the simulated environment and was defined empirically during the performed tests. The purpose of that is to avoid behaviours like an exaggerated acceleration in a very small environment, or slow movement in a large environment.

Controller genes are responsible for 4 behaviors present in our model: rotate right, rotate left, accelerate forward and accelerate backward. The angle to which the robot is facing varies by 15 times the energy of the controller responsible for the rotating behavior: if the controller responsible for turning to the right has an energy of 0.5, the robot will rotate 7.5 degrees to the right (15 *0.5) by position update. The same calculation holds for the rotate left behavior. The robot’s movement speed will vary according to two sensors, the forward and backward acceleration sensors: the speed increases by 7 times the energy of the controller responsible for the forward acceleration, and decreases by 2 times the energy of the controller responsible for the backwards acceleration. In addition, the subject’s maximum frontal speed is 10 units per position update. When moving backwards, the maximum speed is 4 units per update.

### 2.3 Simulated Tasks

In this work, we simulated individuals based on bacteria. Therefore, to assess each individual’s fitness, we chose three tasks with complementary and concurrent relationships among themselves (Table 1), which are inspired by bacterial behavior. The performance achieved in each task was used to evaluate our model’s evolution. The tasks chosen were

1. Objective Orientation, in which the individual must identify and turn his body towards the light,
2. Phototaxy, in which, in addition to identifying where the light is, the individual must move towards it, and finally,
3. Phototaxis with obstacles, in which the individual must avoid threats while performing other tasks.

**Table 1 -.** Classification of relationships between simulated tasks.

| Task 1 | Task 2 | Classification |
| --- | --- | --- |
| Objective Orientation | Phototaxy | Complementary |
| Phototaxy | Phototaxy With Obstacles | Concurrent |
| Phototaxy with Obstacles | Objective Orientation | Concurrent |

The interaction between different tasks results in a decision-making system in living beings defining the priority task at each moment. Tasks that are conflicting - such as “food acquisition” and “protecting offspring” - are defined as concurrent, requiring the individual to eventually stop protecting the offspring in order to go out and look for food. On the other hand, tasks that occur simultaneously or even have a mutually dependent relationship - such as “orientation in the environment” and “food acquisition” - are defined as complementary, since the individual performs both at the same time without one interfering with the other.

In all tasks, the simulated environment is a Cartesian plane, in which the simulated individuals can move freely along the X and Y axes without the presence of walls that impede their passage. All individuals, objectives, and obstacles are generated within the limits −100 to 100, both on the X-axis and the Y-axis. Each performed simulation occurred in 800 individual position updates, and their interaction with the environment was evaluated differently for each task. To avoid false positives in our population, the same individual participated multiple times in the same task; its fitness is considered the average of all simulations performed. The more complex the task, the more simulations were needed to eliminate false positives.

In order to evaluate the ideal amount of simulations to reduce false positives without compromising the model’s efficiency, we carried out a study using the “phototaxy” task (Figure 4). In this study, we evaluated three populations: the first had low fitness, not being proficient in the task, and had a high chance of false positives. The second was composed of individuals who have already trained enough to have average fitness but still have false positives. Finally, a population with high fitness was evaluated, in which all individuals always performed very well on the task, with a low rate of false positives. For this study, each individual in the population was evaluated in an increasing number of simulations, with the calculated fitness being compared to the final fitness of the individual in question.

**Figure 4:**
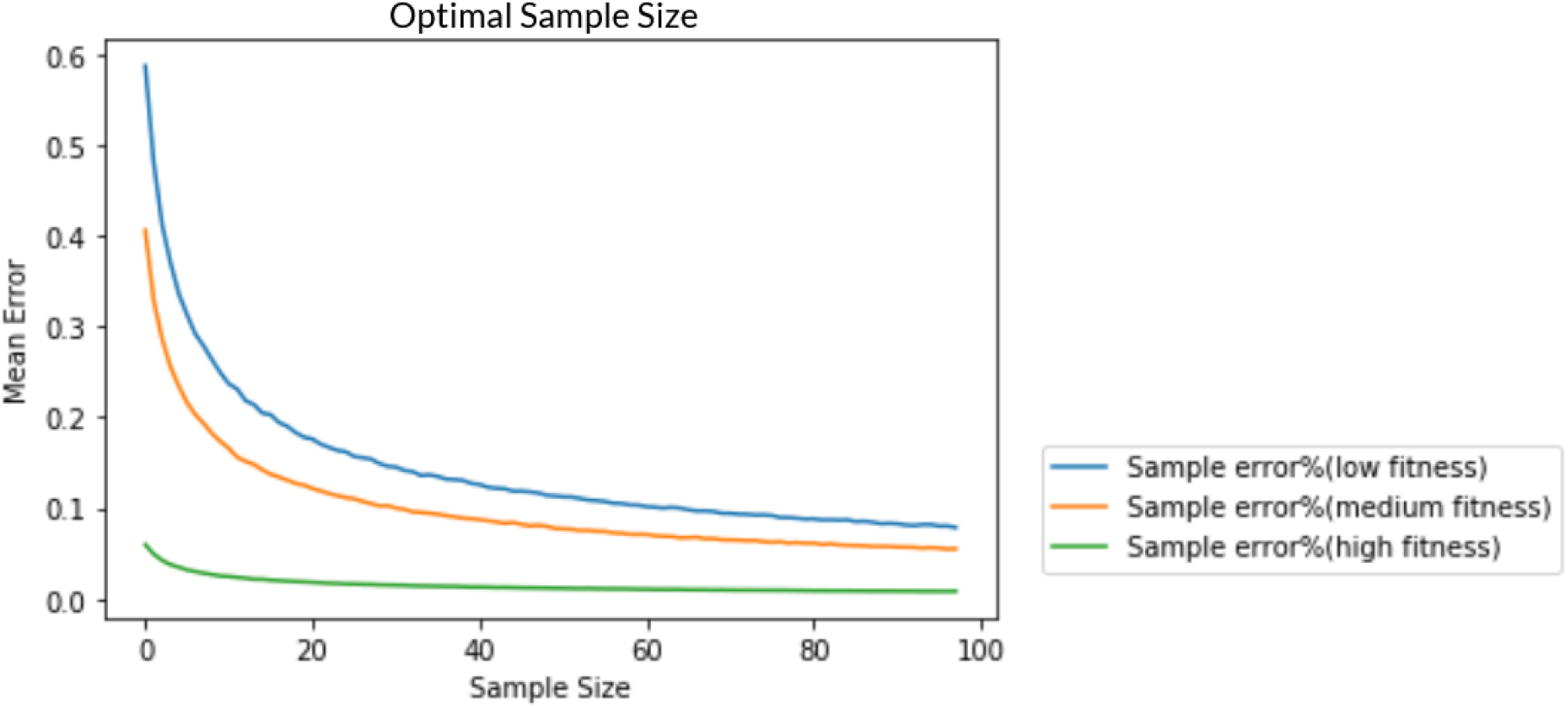
Evaluation of the fitness error rate when different numbers of simulations occurred for different populations. In blue, a population with low average fitness. In yellow, a population with good average fitness. In green, a population where average fitness is optimal. On the X axis, we have the number of simulations performed for the assessment of each individual’s fitness, and on the Y axis, we have the mean percentage of error between the actual fitness and that attributed after the simulations.

#### 2.3.1 Objective Orientation

In Objective Orientation simulations, the individual is positioned in the center of the simulated environment and must be able to orient its vision towards lights that will appear in the environment. When the individual is successfully able to look at a light, it disappears and another light is created at a random location in the environment. The individual’s fitness is composed of the amount of lights that he was able to orient himself to. For the elimination of false positives, the fitness results from the average of 5 simulations in this task. For the individuals physical structure, 7 proximity sensors are used and react to the lights that appear. In addition, the 2 controllers responsible for turning the body to the right and left are also used.

#### 2.3.2 Phototaxis

For Phototaxis simulations, the individual is positioned at a random location in the simulated environment and must be able to move its body until it gets close to lights that will appear in the environment, as seen in Figure 5. A light is considered captured when the individual is at least 8 units of distance from the light. When a light is caught, it disappears and another light is created at a random location in the environment. The individual’s fitness is composed of the amount of lights that he was able to capture. For the elimination of false positives, the fitness results from the average of 10 simulations in this task.

**Figure 5:**
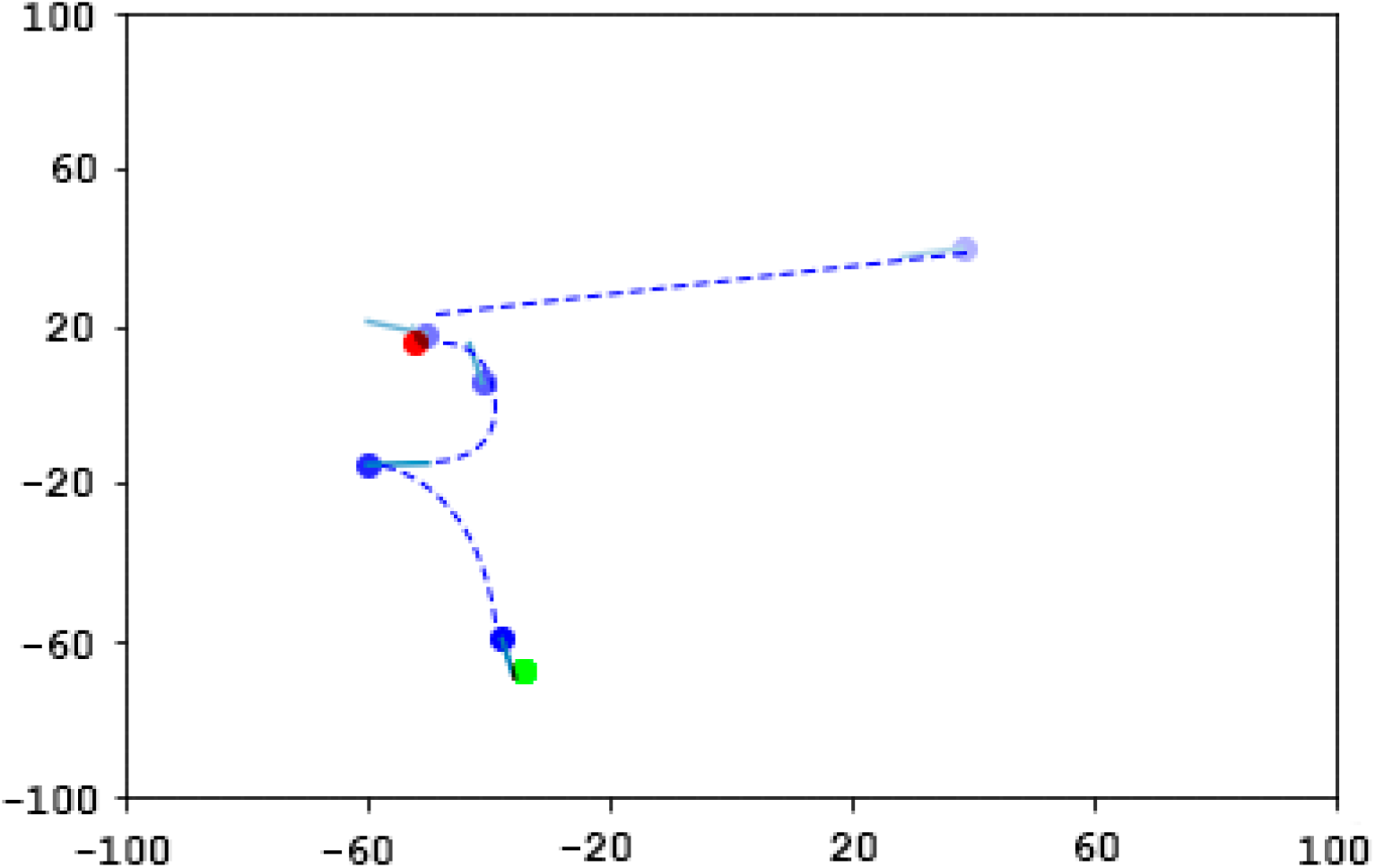
Example of a simulated individual during the “Phototaxy” task. In blue, the simulated individual with a line indicating where he is facing. In red and green, the first and second objectives captured, respectively. The path taken by the individual is represented by the blue dotted line.

The individuals’ physical structure in this activity is composed of 7 proximity sensors (the same ones used in the “Goal orientation” task) and 4 controllers: 2 responsible for turning the body to the right and left and 2 that control the action of accelerating forward and backwards.

#### 2.3.3 Phototaxis with Obstacles

Phototaxis with Obstacles simulations are very similar to common phototaxy. In this task, the bacteria’s objective is to capture lights that appear in the environment, as seen in Figure 6. However, 4 obstacles are added to the environment: if the individual comes into contact with such obstacles, reaching less than 15 units away from them, will receive a penalty on its adaptability and the obstacle will be moved to a random spot in the environment. The individual’s fitness, in this case, is composed of the amount of lights he managed to capture minus the amount of contacts with obstacles. For the elimination of false positives, the fitness results from the average of 15 simulations in this task.

**Figure 6:**
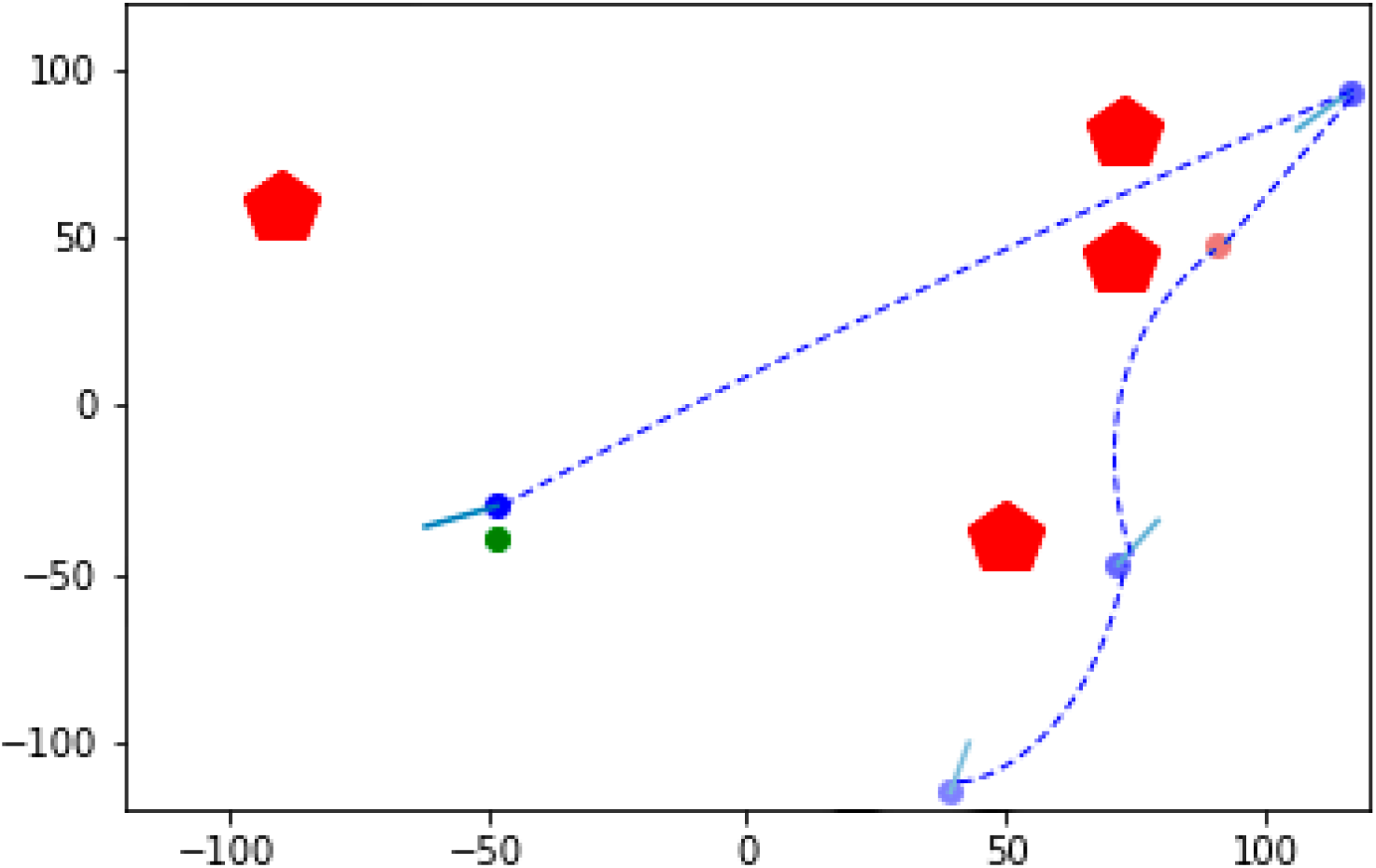
Example of a simulated individual during the task “Phototaxis with Obstacles”. In blue, the simulated individual with a line indicating where he is facing. In red and green, the first and second objectives captured respectively. The red pentagons represent the obstacles to be avoided, there was no clash with obstacles in this simulation. The path taken by the individual is represented by the blue dotted line.

For the physical structure of individuals, we kept the same controller structure as in the “Phototaxy’’ task. Regarding the sensors, in addition to the 7 proximity sensors, we also use the frontal sensor responsible for differentiating between obstacle and objective.

### 2.4 Evolutionary Algorithm

The created networks were optimized through an evolutionary algorithm, being evaluated in the tasks defined above. After selecting one of the tasks, an initial population of 100 random individuals is created with physical structure equivalent to the task. Each of them then participates in simulations that generate individual fitnesses. The next generation is composed of: 10 individuals who had the greatest fitness in the previous generation, 80 offspring formed by sexual reproduction (crossing-over) and 10 individuals coming from gene flow (immigration). The simulated evolutionary process can be seen in Figure 7.

**Figure 7:**
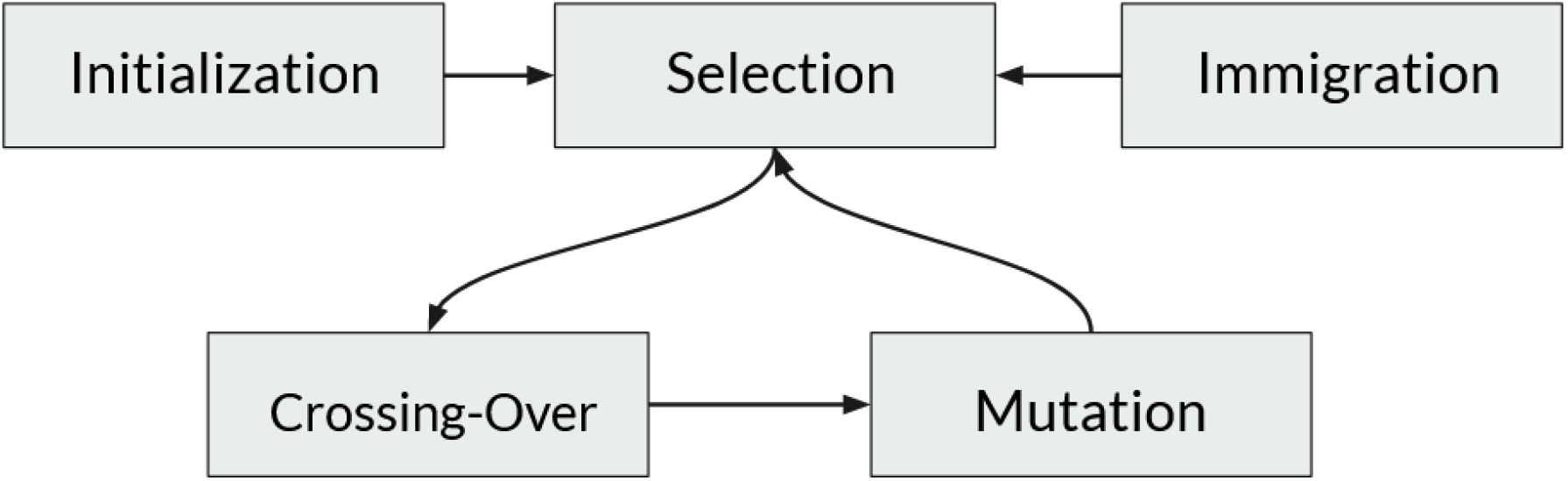
Flowchart of the simulated evolution process.

To perform sexual reproduction with crossing-over, two surviving parents are chosen randomly, and each edge (relationship between two genes) of the child graph has a 50% chance of being copied from each parent. After the vertices are copied, 0.1% of them are mutated, adding genetic variability to the offspring. During the mutation process, the edges are chosen randomly, with the network having a 50% chance of deactivating this relationship between two genes, and a 50% chance of replacing it with a random value between −1 and 1. There is, thus, the representation of a newly created inhibition or excitation relationship. To carry out the gene flow event (immigration), 10 new individuals based on the survivors (parents) are added to the population and have 5% of their edges mutated. This process is important to increase the genetic variability of the population.

After the creation of individuals through sexual reproduction and gene flow, a new generation is started, initiating a new evaluation of individuals, death of individuals with low fitness and generation of new children and “immigrants”. The number of generations required for the model to reach a local maximum is linked to the complexity of the evaluated task. For the “Goal Orientation” task, 10 generations are needed to reach a development plateau. On “Phototaxy”, 40 generations are needed. Since “Phototaxy with Obstacles” is a task with greater complexity, 60 generations were needed to identify the local maximum.

A summary of the simulated parameters in each task can be seen in Table 2.

**Table 2 -.** Summary of used parameters in each simulated task. PS: proximity sensors; DS: differentiation sensor; RC: rotation controllers; AC: Acceleration controllers.

| Simulated Task | PS | DS | RC | AC | Number of Simulations | Fitness Calculation | # of Generations |
| --- | --- | --- | --- | --- | --- | --- | --- |
| Objective Orientation | Yes | No | Yes | No | 5 | # of lights seen | 10 |
| Phototaxy | Yes | No | Yes | Yes | 10 | # of lights captured | 40 |
| Phototaxy With Obstacles | Yes | Yes | Yes | Yes | 15 | # of lights captured minus # of obstacle collisions | 60 |

## 3. RESULTS

### 3.1 Network Size Analysis

Our objective was to evaluate how biologically inspired topological changes influence the learning process in an AGRN, but first, we needed to evaluate the control behavior of the network when working with different amounts of genes.

Figure 8 shows the mean fitness of the population using different net sizes after training, assessed in the phototaxy task. To increase the amount of genes, only processor nodes were inserted, as we are not modifying the structure of sensors and controllers of individuals. The confidence interval for this study, as well as for all subsequent ones, was calculated using the bootstrap method (EFRON; TIBSHIRANI, 1994).

**Figure 8:**
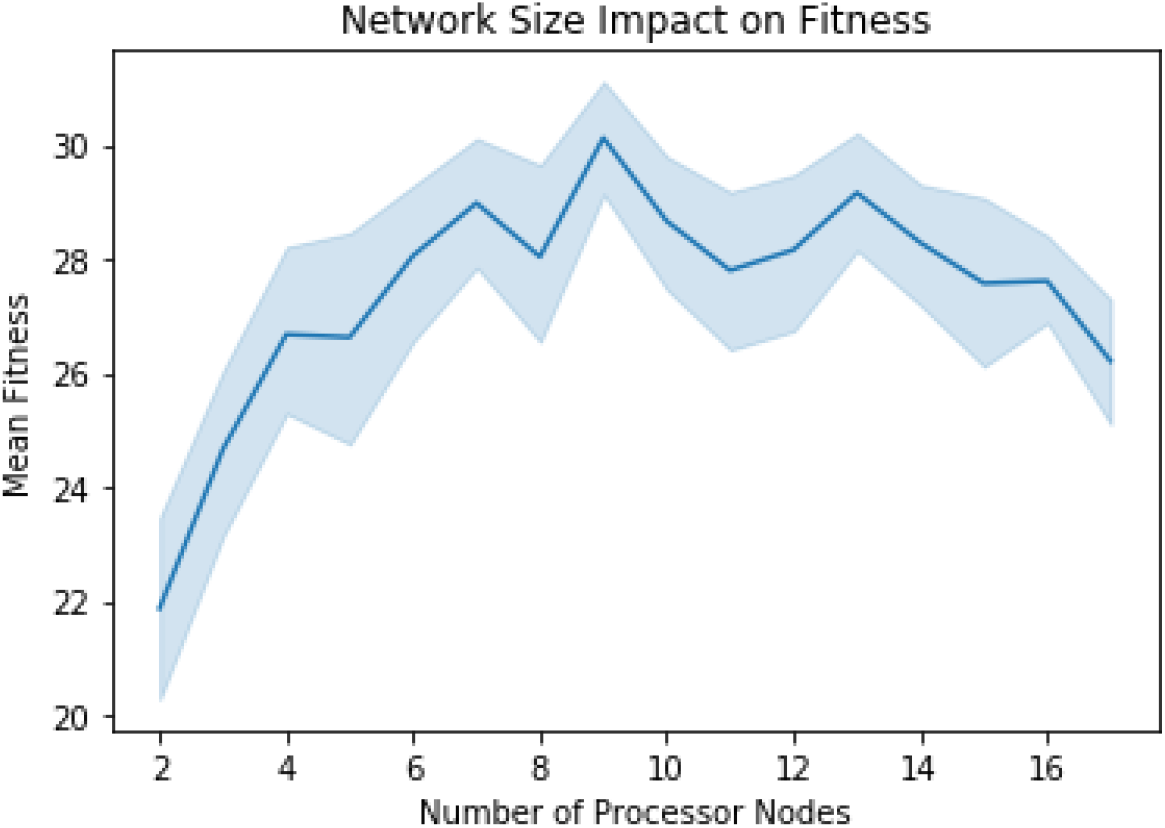
Evaluation of the average performance of the post-training populations for 32 simulations (confidence intervals of 68%). Networks with a number of processor nodes varying between 2 and 18 were used.

This analysis shows that a greater amount of genes is not necessarily reflected in a better fitness. Due to the random nature of the network, excessively large amounts of genes hinder task learning. The larger the network, the less likely a controller is to be excited by a relevant sensor, and the greater the amount of noise between sensors and controllers.

### 3.2 Mutation Rate Analysis

The mutation rate (chance of a gene being modified during the simulation) influences the learning of an AGRN. Since we are initially working with a much smaller number of genes than most animals, we need to identify the need or not for a variable mutation rate for different network sizes. Figure 9 shows the average fitness after training on the phototaxy task for different mutation rate values. The mutations performed to simulate the gene flow were proportionally altered. The networks used are composed of 7 sensor nodes and 4 controllers, resulting in 11 static nodes in the network, in addition to a variable number of processor nodes. In the cases observed, networks with 4 and 9 processor nodes were used, totaling 15 and 20 genes, respectively. The networks keep the same number of nodes throughout the simulation.

**Figure 9:**
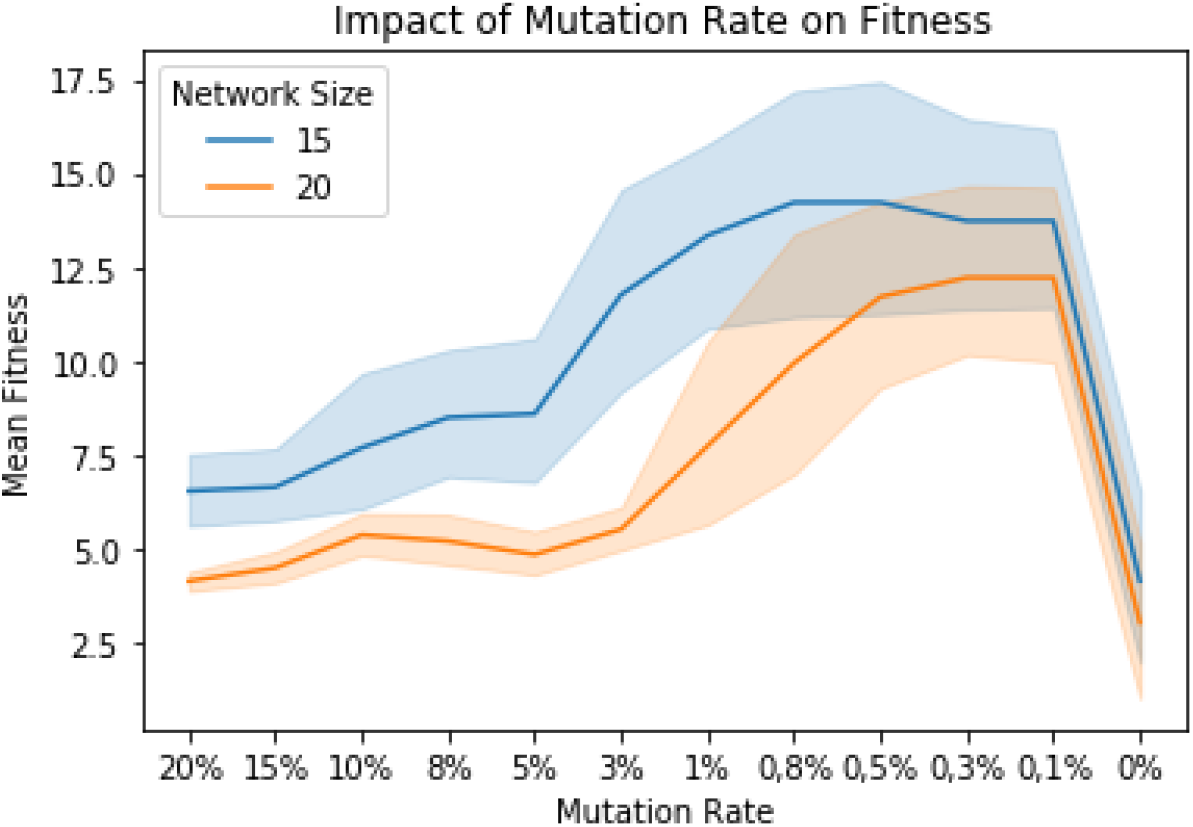
Average performance evaluation for 32 simulations, when different mutation rates are used (68% confidence intervals). Mutation rates range from 20% to 0%. In blue, populations with individuals with 15 genes in their network. In orange, populations with individuals with 20 genes in their network.

As seen, regardless of net size, low mutation rates result in greater success while training an AGRN. Despite this, by completely removing the mutation rate from the simulations, recombination becomes the only way to modify the network for the children generated during the simulations, making it impossible to improve the individuals in the evaluated tasks. This happens because, without mutation, there is no generation of new network interactions, removing gene diversity in the population.

### 3.3 Multiple Tasks Learning

The next target of study was the impact of the gradual increase in the complexity of tasks on the population’s ability to evolve. Let us consider that species did not appear from nowhere, rudimentary life forms evolved to more complex ones (ALBERTS et al., 2017). To represent this, in Figure 10, we compare the learning ability of randomly initiated populations with populations that have been trained in less complex tasks.

**Figure 10:**
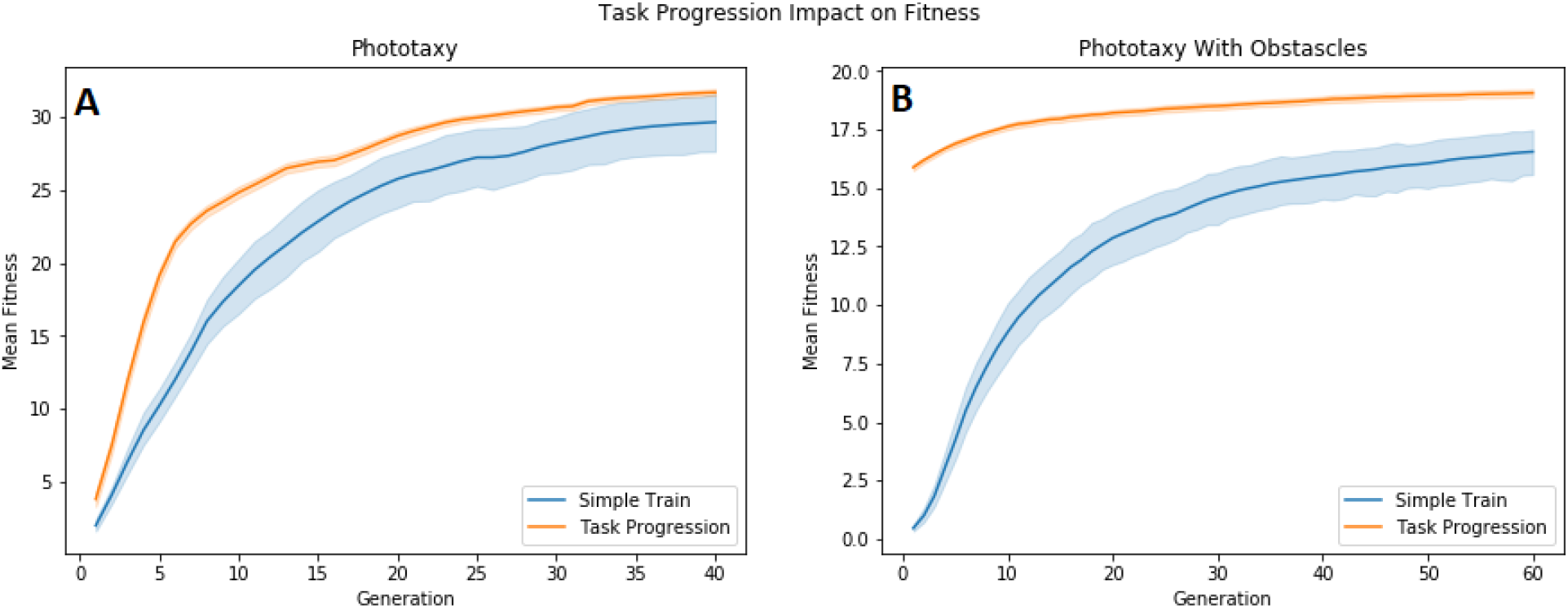
Average performance Evaluation of the network for 32 simulations (confidence intervals of 68%) in the tasks: “Phototaxy” and “Phototaxy with obstacles’’, for two different strategies. In blue, naive populations (no previous training). In orange, populations that had previous training in less complex tasks, “Goal Orientation” for Figure A and “Phototaxy” for Figure B.

In the case of phototaxy, the physical structure used was the basic structure of the task (7 sensor genes and 4 controllers), plus 8 processor genes. The population was pre-trained in the “Objective Orientation” task.

For “Phototaxy with Obstacles”, the object differentiation sensor was added, totaling 8 sensor genes and 4 controllers (8 processor genes were used). Individuals from these populations were pre-trained in the “Phototaxy”task.

It is observed that prior training in less complex tasks not only increases the maximum fitness a population can achieve, but also creates extremely stable populations, that is, children created over generations are consistently able to perform as well as their parents, since the networks present in the population are more stable. In populations that started training in more complex tasks, a small portion of the population achieves higher fitness, but has difficulty passing this information on to their descendants. This is because during reproduction and mutation, their less stable networks end up losing important information, resulting in children with little chance of survival.

### 3.4 Network Growth Events

One of the natural events that help prevent loss of function due to mutation of specific genes is gene redundancy, that is, the existence of multiple copies of the same gene found in the genome. These copies ensure the presence of vital proteins and create the possibility for the generation of new genes arising from the mutation of functional genes (ALBERTS et al., 2017). Based on this, Figure 11 shows the comparison of two network growth events: single gene duplication and whole genome duplication. For both events, the population was evaluated in the phototaxis task, and started with random populations (they were never trained in other tasks). The physical structure used was the basic structure of the task (7 sensors and 4 controllers). In addition, each individual started the simulations with 8 processor genes and, after 80 simulated generations, each individual ended up with 16 processor genes, due to the network growth events that occurred. In whole genome duplication, the event takes place in generation 40. During this event, each individual in the population has all of its processor genes duplicated, as well as the interactions between them. For the study of single gene duplication, 8 events occur during generations 8, 16, 24, 32, 40, 48, 56, and 64. A random processor gene was duplicated in each of these events.

**Figure 11:**
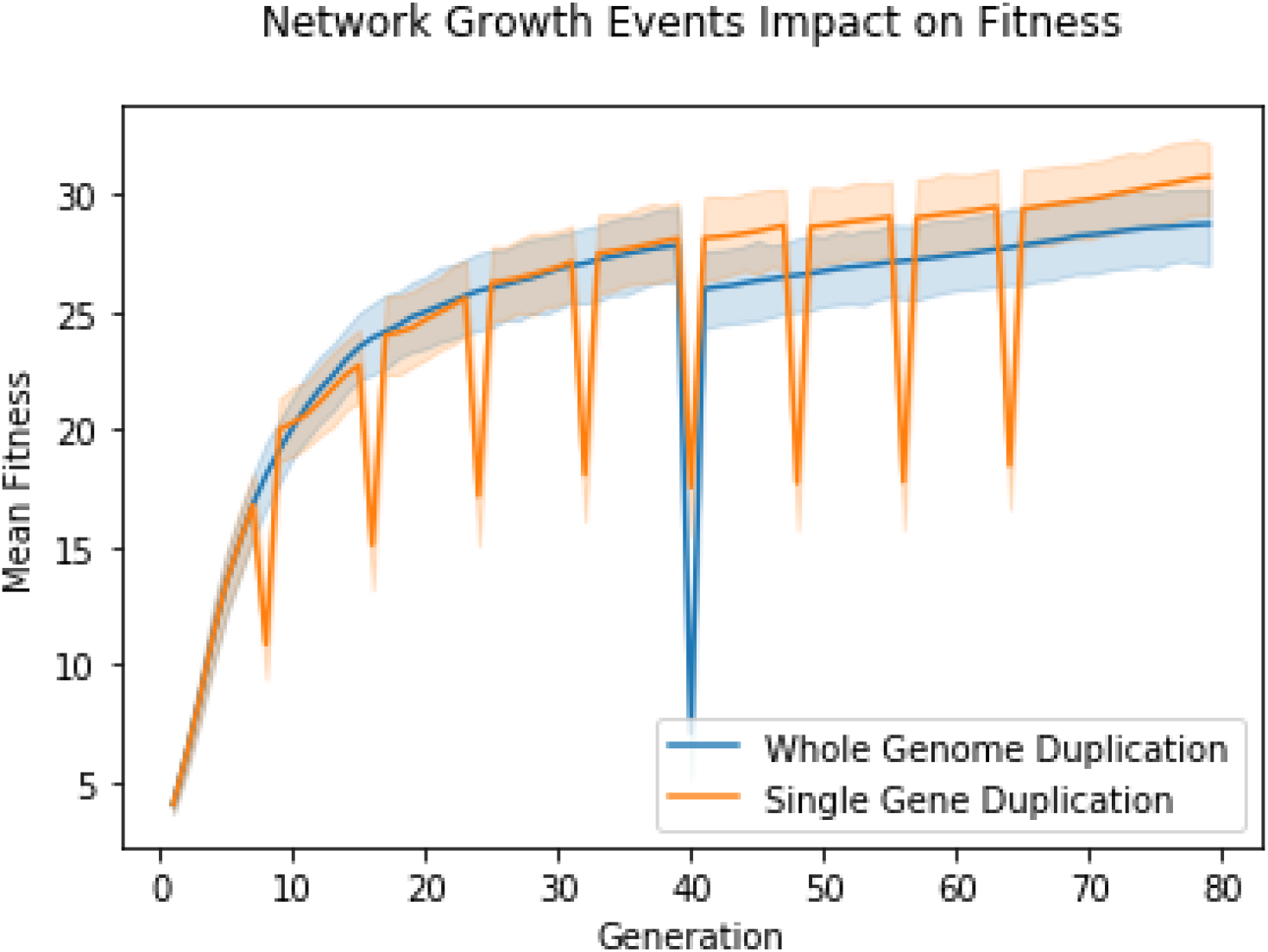
Assessment of the impact of network growth events on average population fitness for 32 simulations (68% confidence intervals). In blue, average fitness of populations that experienced full genome duplication events in generation 40. In orange, populations that experienced 8 single gene duplication events in generations 8, 16, 24, 32, 40, 48, 56, and 64.

Due to the random nature of the model, network growth events, while being necessary for the inclusion of more complex tasks, initially decrease the average fitness of the population by having the chance to duplicate genes that do not play an important role, adding noise to the network. Thus, this study, by duplicating the entire genome, despite ensuring the duplication of all functional genes, ends up simultaneously duplicating all the noises that are already present in the network. Consequently, it ends up causing a delay in learning of the model. Single gene duplication turned out to be a much more efficient network growth event. When suffering small disturbances, the model is able to re-stabilize its network and resume learning without harm.

### 3.5 Gene Stress Simulation

Developing well-defined models and tasks enables simulation of hypothetical situations that would not be well observed or even impossible to analyze *in vivo* or *in vitro*. In this research, we simulated a population of bacteria being targeted by hypothetical drugs that attack specific physical characteristics of bacteria, more precisely, the sensing capabilities, represented by the network sensor genes, or motor, represented by the controller genes.

For this, we started our simulation with the 10 best individuals from a population trained in the “Phototaxy” task. We then added gradual mutations to random individuals in the population, always in vertices (relationships between two genes) related to sensor or controller genes.

Sensor genes have a bottleneck position when inputting any information into the simulated network. Therefore, changing them interferes with the way the entire network processes the environmental information received. This can be seen in the near incapacitation of the population, as seen in Figure 12, when hypothetical drugs specifically target sensor genes.

**Figure 12:**
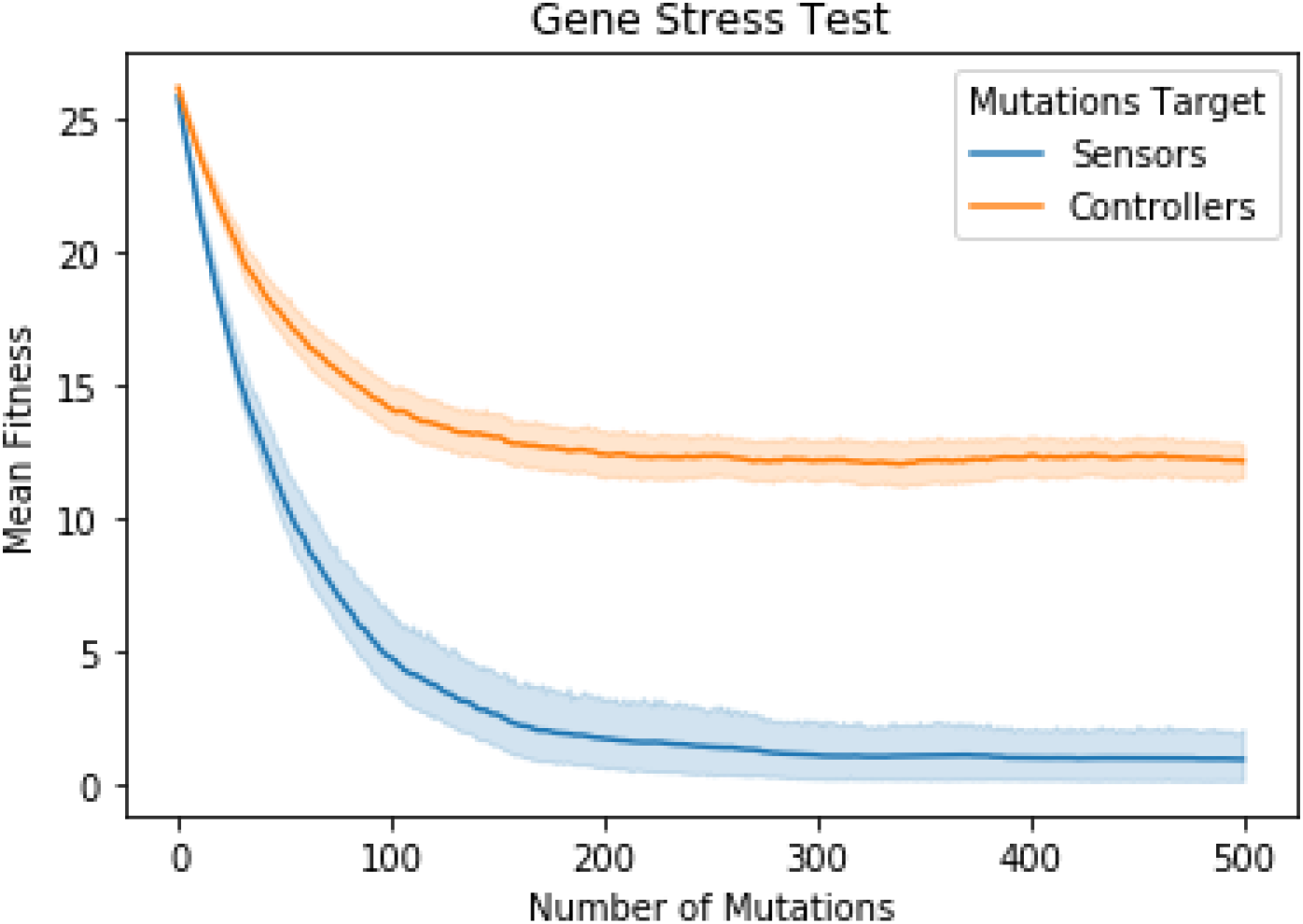
Assessment of the impact of sensor- and controller-directed mutations on the population’s mean fitness (confidence intervals of 68%). In blue, average fitness of 32 simulations in which a population undergoes targeted mutations to its individuals’ sensor genes. In orange, average fitness of 32 simulations in which a population undergoes mutations aimed at controller genes of its individuals.

Controller genes, in turn, are positioned in the middle of the AGRN, and although mutations in their interactions decrease the quality of the evaluated task, the dynamics of the network is maintained. This results in a final population that is still capable of performing the task, but inefficiently.

## 4. CONCLUSION

In this work, we used AGRNs to simulate evolution, identifying how environmental variation events (represented through tasks) and genome growth affect the development of the evaluated populations. The results indicate that a gradual increase in the complexity of the performed tasks is beneficial for the evolution of the model. We have also seen that larger gene regulatory networks are needed for more complex tasks, with single gene duplication being a good evolutionary strategy for growing these networks, as opposed to whole genome duplication.

From this perspective, the investigation carried out shows that further studies of processes and phenomena involving GRNs reveal not only ways to optimize the use of AGRNs, but also provide insights into aspects and events that influenced the development of life on Earth.

## Footnotes

1 Source: <https://cyberbotics.com/doc/guide/epuck>. Acessed in: 20 ago. 2021.

